# Dual oxidase gene *Duox* and Toll-like receptor 3 gene *TLR3* in the Toll pathway suppress zoonotic pathogens through regulating the intestinal bacterial community homeostasis in *Hermetia illucens* L.

**DOI:** 10.1101/844696

**Authors:** Yaqing Huang, Yongqiang Yu, Shuai Zhan, Jeffery K. Tomberlin, Dian Huang, Minmin Cai, Longyu Zheng, Ziniu Yu, Jibin Zhang

**Affiliations:** State Key Laboratory of Agricultural Microbiology, National Engineering Research Center of Microbial Pesticides, College of Life Science and Technology, Huazhong Agricultural University, Wuhan, P.R. China 430070; Institute of Plant Physiology & Ecology, SIBS, CAS, China 200032; Department of Entomology, Texas A&M University, USA

**Keywords:** *Hermetia illucens*, zoonotic pathogens, *BsfDuox*, *BsfTLR3*, gut microbiome.

## Abstract

Black soldier fly (BSF; *Hermetia illucens* L.) larvae can convert fresh pig manure into protein and fat-rich biomass, which can then be used as livestock feed. Currently, it is the only insect approved for such purposes in Europe, Canada, and the USA. Pig manure is rich in zoonotic pathogens (e.g., *Staphylococcus aureus* and *Salmonella* spp.). BSF larvae inhibit these zoonotic pathogens; however, the mechanism is unclear. We employed RNAi, qRT-PCR, and Illumina MiSeq bacterial 16S rDNA high-throughput sequencing molecular techniques to study the interaction between the two immune genes (*Duox*in Duox-reactive oxygen species [ROS] immune system and *TLR3* in the Toll signaling pathway) and zoonotic pathogens to determine the mechanisms resulting in pathogen suppression.

Results indicated that *Bsf Duox-TLR3* RNAi increased bacterial load but decreased the relative abundance of *Providencia*and *Dysgonomonas*intestinal symbionts. Concurrently, *Bsf Duox-TLR3* RNAi inactivated the NF-κ B signaling pathway, downregulated the expression of antimicrobial peptides, and diminished inhibitory effects on zoonotic pathogen. The resulting dysbiosis stimulated an immune response by activating *BsfDuox* and promoting ROS, which regulated the composition and structure of the gut bacterial community.

Thus, *BsfDuox* and *BsfTLR3* are important factors in regulating the gut key bacteria *Providencia*and *Dysgonomonas* homeostasis while inhibiting target zoonotic pathogens.

## Introduction

*Hermetia illucens*L. (Diptera: Stratiomyidae) is a saprophytic insect whose larvae consume various organic wastes of environmental concern and convert them into biomass rich in protein and fat[1]. BSF larvae (BSFL) provide environmental aid and inhibit zoonotic pathogen loads in livestock wastes, such as pig manure. Meanwhile, Liu et al. (2008)[2] determined that BSFL can reduce *E. coli* in dairy manure. Furthermore, Lalander et al. (2015) [3]discovered that BSFL not only reduce *Salmonella* spp. but also reduce viruses in organic wastes.

Several researchers have explored how BSFL inhibit these zoonotic pathogens. Park et al. (2015)[4] characterized an *H.illucens* defensin-like peptide which have activity against gram-positive bacteria. Elhag et al. (2017)[5] identified seven gene fragments responsible for the production of three types of antimicrobial peptides. Zdybicka-Barabas et al. (2017)[6] examined *H.Illucens* hemocytes and found *E coli*-challenged larvae increased phenol oxidase, lysozyme and anti-gram-positive bacterium activity.

To combat infection, insect relies on innate defense reactions to defense pathogens by producing antimicrobial peptides, phenoloxidase and H_2_O_2_.In *Drosophila* (Diptera: Drosophilidae), the Toll signaling pathway is mainly induced by gram-positive bacteria and fungi[7]. The recognition receptors *GNBP3* and *PGRP-SA* in hemolymph are combined with β-1,3-glucoside on the surface of fungal cells and lysine-type peptidoglycan on gram-positive bacteria cells. The Toll-like receptor pathway ligand Spatzle is cleaved and proteolysed, and it activates the nucleic acid transcription factors *Dif* and *Dorsal* to mediate antibacterial peptide gene expression.

Toll-like receptors (TLRs) are a class of type I membrane receptors with an extracellular amino terminus and a conserved cytoplasmic region. TLRs, which widely exist in antigen-presenting cells, are a pattern receptor involved in recognizing molecular structures (e.g., PAMPs) specific for microbial pathogens and have an important effect on innate and adaptive immune response. With routine microbial burdens, such as those found in the absence of infection, the Toll pathway at low activation levels. However, acute pathogenic bacterial infection transiently increases nuclear factor kappa B (NF-κB)-dependent innate immune signaling.

The insect gut immune system produces microbicidal ROS by dual oxidase (Duox) to restrict the proliferation of invading microorganisms. In addition, ROS is involved in regulating the healing process of intestinal trauma and also functions as a signaling molecule to initiate other self-balancing signaling pathways[8]. The intestinal bacterial community is associated with host immunity and bacteriostasis. Microbial flora modulate anti*-*pathogen effects of some immune genes plausibly through activating basal immunity[9]. For example, in *Bactrocera dorsalis*, ROS which is induced by *BdDuox* gene plays a key role in intestinal bacterial community homeostasis[10]. ROS is an important immune mechanism for many insects to protect themselves against pathogenic microorganisms, such as bacteria and fungi. Kumar et al. (2010) [11] and Mitochondria in *Anopheles gambiae* Giles (Diptera: Culicidae) intestinal cells and *Enterobacter* of intestinal bacteria can also increase ROS levels to decrease the infection rate of the malarial parasite *Plasmodium*[12].

The Duox regulatory pathway also contributes in maintaining gut–microbe homeostasis in insects[13]. Gut membrane-associated proteins Meshregulate *Duox* expression throughan arrestin-mediated MAPK/JNK/ERK phosphorylation cascade and play an important role in controlling the proliferation of gut bacteria. Expression of both *Mesh* and *Duox*is correlated with the gut bacterial microbiome[14].

Recent surveys of black soldier fly gut microbiota revealed a diverse community dominated by Bacteroidetes and Proteobacteria[15]. The native gut microbiota (or indigenous microbiota) mediates novel protection function such as pathogen cleareance[16, 17]. For example, the microbial community of the red flour beetle *Tribolium castaneum* Herbst(Coleoptera:Tenebrionidae) offers protection against *Bacillus thuringiensis* bv *tenebrionis*[18]. Experiments on silkworm *Bombyxmori* L.(Lepidoptera: Bombycidae) demonstrated that lactic acid bacteria in the gut enhance host resistance against *Pseudomonas aeruginosa*. Despite the involvement of native gut microbiota in combating infections, the manner by which the immune system of BSFL regulates gut microbiota homeostasis to suppress zoonotic pathogens remains unknown.

In this study, we examined the antimicrobial activity of immune genes BSFL dual oxidase (*BsfDuox*) and TLR 3 (*BsfTLR3*), which represent two classic immune pathways. We explored the (1) reduction in pathogen loads in pig manure by BSFL, (2) expression profile of immune genes *BsfDuox* and *BsfTLR3* in BSFL, (3) expression profile of immune genes after oral pathogenic bacterial challenge, (4) whether BSFL reduces suppression on zoonotic pathogens after *BsfDuox-TLR3* RNA interference (RNAi), and (5) the dynamic change in intestinal bacterial community after RNAi of *Duox-TLR3* genes and relationship with the suppression of zoonotic pathogens.

## Material and Methods

### Rearing and dissection of *H. illucens L*

A colony of BSF (Wuhan strain) used in this study was taken from the State Key Laboratory of Agricultural Microbiology of HZAU. *H. illucens* larvae were reared for 10 days at 27 °C and 70%-80% relative humidity on artificially sterilized feed (75 g of bran, 75 g of corn flour, and 350 g of water). Third instars were surface-sterilized. After 70 % of the ethanol washed once, it is washed three times in sterile water. Approximately 60 larvae from each experiment were dissected in a sterile distilled water with a sterilized tweezers under a stereo microscope[19]. The dissected guts were transferred to a sterile PBS test tube and homogenized. The homogenate was used for different experiments.

### Zoonotic pathogenic bacteria assay in pig manure conversion

Surface-sterilized BSFL at 8-10 days old were placed into 100 g of fresh pig manure collected from a facility located near the university. Conversely, the control group consisted of equal amounts of sterile distilled water introduced into another set of fresh pig manure. All treatments were repeated three times. The detection method of zoonotic pathogens was based on the national standards of China, in which *Staphylococcus aureus* was detected[20] The method is described as follows. The selected medium for *S. aureus* was prepared by placing 6.3 g of Baird-Parker agar in 95 ml of distilled water; heated to boil until completely dissolved, autoclaved at 121 °C for 15 min, and agitated well after sterilization, thereby preventing agar from depositing at the bottom and solidifying. The medium was then cooled to 50 °C, added with 5 ml of potassium citrate-potassium yolk enrichment solution, gently agitated, and poured into a plate. To count *S. aureus* colonies, we first collected manure samples (5 g) at different times after the larvae were inoculated (0, 2, 4, 6, and 8 days). We mixed the samples with 45 ml of sterile physiological saline in a 100 ml sterile conical flask and agitated for 16 min at 180 rpm. The sample was placed at room temperature (RT) for 5 minutes, and the upper layer was withdrawn for subsequent tests. Dilutions were prepared using normal saline, and 0.1 ml of each dilution was mixed with the appropriate select medium and plated. Pathogen counts were determined using a selective enrichment method on these plates, following incubation of plates at 37 °C for 48 h. *S. aureus* colonies exhibited a diameter of 2-3 mm, gray to black appearance, light-colored edge, turbid zone, and transparent ring on the outer layer. The number of average *S. aureus* of three plates was expressed as CFU/g (Standards Press of China, 2010)

Meanwhile, *Salmonella* spp. were detected by GB 4789.4-2016[21]. The medium for *Salmonella* spp. was prepared by adding 5 g of bismuth sulfite agar to 100 ml of distilled water, stirred and boiled for 1 min, cooled to 50 °C-55 °C, plated, and used overnight. *Salmonella* spp. colonies were counted by extracting manure samples (5 g) at different times after larval inoculation (0, 2, 4, 6, and 8 days), collected, mixed with 45 ml of sterile physiological saline in a sterile 100 ml conical flask, and agitated for 15 min at 180 rpm. The sample was undisturbed for 5 min at RT, and the upper layer was withdrawn for subsequent tests. Dilutions were prepared using normal saline. Approximately 0.1 ml of each dilution was mixed with the appropriate selective medium and then plated. Pathogen counts were determined using a selective enrichment method on these plates, following the incubation of plates at 37 °C for 48 h. *Salmonella* spp. colonies were black with a metallic luster, tan or gray, and the medium surrounding the colonies may be black or brown; some strains formed gray-green colonies, and the surrounding medium remained unchanged. The number of average *Salmonella* spp. of the three plates was expressed as CFU/g (Standards Press of China, 2016).

### Total RNA isolation and cDNA synthesis

The dissected gut samples were used for RNA isolation. Total RNA was extracted from different developmental stages of BSF, including egg, first-fourth instars, and adults after mating (13-15 days after eclosion) by using TRIzol extraction kit (Invitrogen USA). In addition, total RNA was isolated from RNAi-*Duox-TLR3* and RNAi-*egfp* groups. The experiments were performed three times. RNA quality was analyzed by NanoDrop 2000 spectrophotometry (ThermoFisher Scientific Inc., Waltham, MA, USA) at 260 nm. 1 μg of total RNA was reversely transcribed into first-strand cDNA by using synthesis kit (Vazyme).

### Sequence analysis of full-length *BsfDuox* and *BsfTLR3*

The genome of BSF was sequenced and assembled. The genome size of BSF was 1.3 G, which is the largest among the Diptera that has been sequenced (data not published). We searched the *BsfDuox* and *BsfTLR3* genes in the BSF genome database (the URL has not been publicly available) by using the Duox protein sequence and TLR3 protein sequence of the fruit fly. The highest similarity sequence of protein was the protein sequence of Duox and TLR3 of BSF. Specific primers for *BsfDuox* and *BsfTLR3* were designed using Premier5.0 software. The transmembrane domains in *BsfDuox* and *BsfTLR3* were identified using TMHMM online software (http://www.cbs.dtu.dk/services/TMHMM-2.0/), and the structural domains of *BsfDuox* and *BsfTLR3* were predicted using the simple modular architectural research tool (SMART, version 7.0, http://smart.embl-heidelberg.de/).

### Microbial oral infection

Third-instar larvae (8 days old) were fed with an artificial diet containing concentrated microbe solution (1×10^8^ CFU/ml), 5% sucrose solution only served as the control.

Bacteria used for oral infection were grown in lysogeny broth medium at 28 °C and 180 rpm. Exponential microbial culture (OD_600_=1.0) was used for all the experiments as previously described [22]and then centrifuged for 15 s (8,000 g) with aseptic distilled water. The resulting bacterial counts in each sample were adjusted to 1×10^8^ CFU/ml by aseptic distilled water. The larvae fed with a 5% sucrose diet only served as the control. For the analyses of *BsfDuox* and *BsfTLR3* gene expression and ROS level changes after oral infection, the gut samples of different treatments were collected at different times post-oral infection(POI). The microorganisms used in this study were pathogens *S. aureus* and *Salmonella* spp.

### Real-time quantitative PCR (qRT-PCR) analysis

In all cases of gene expression analysis, three third-instar larvae were collected for RNA extraction and cDNA synthesis. qRT-PCR was performed using a Bio-Rad CFX system (Bio-Rad, Hercules, CA, USA) with a 384-well plate. Each PCR mixture consisted of 7.8 μl of SYBR Green Mix (Vazyme), 10 nM of each primer, and 2 μl of cDNA (diluted 1:10). The amplification program was consist of pre-incubation at 95 °C for 3 min, followed by 39 cycles of denaturation at 95 °C for 20 s, annealing at 56 °C for 20 s, and extension at 72 °C for 20 s. After the fluorescence quantitative PCR was over, the dissolution curve was analyzed to ensure specific amplification by a 0.5 °C increase for 5 sin each cycle started with 65 °C for 10 s until 95 °C. Real-time fluorescence quantitative PCR results were measured by 2^-ΔΔCt^ method as described previously [23]. All the samples were analyzed in triplicate, and the levels of the detected mRNA determined by cycling threshold analysis were normalized using β-actin as the control. The primer pairs used in qRT-PCR analysis are shown in Supplemental Table 1. The loads of total bacteria were quantified by qRT-PCR using 16S RNA gene-specific primers [24]and normalized by using β-actin as the control via a previously described method [25].

### Measurement of total in vivo ROS

The intestines of individual third-instar larva were rapidly hand-dissected in PBS. The dissected intestines were ground with PBS. The quantification of ROS was completed according to the corresponding kit instructions provided by the Institute of Nanjing Jiancheng Bioengineering. In brief, we weighed the tissue accurately, added nine times volume of normal saline, centrifuged the sample at 5,000 g for 5 min, and measured the supernatant. The absorbance values of each tube were measured under 405 nm by using UV756 (756 spectrophotometer, Shanghai Opal Instrument Co., Ltd.). The protein concentration of each tube was determined by NanoDrop2000 (ThermoFisher Scientific Inc., Waltham, MA, USA). The ROS value of each tube was calculated using the following formula: ROS concentration = (measured OD value−blank OD value) × 163 / (Standard OD value−blank OD value) × sample protein concentration.

### Double-stranded RNA (dsRNA) synthesis and delivery by injection

The *BsfDuox* and *BsfTLR3* sequence fragments were amplified by normal PCR with premier 5.0 to design of specific primers conjugated with the T7 RNA polymerase promoter. The primer pairs used in dsRNA synthesis are shown in Supplemental Table 1.1 μg PCR product was used as the template for dsRNA synthesis utilizing the T7 Ribomax Express RNAi System (Promega, Madison, WI, USA). The dsRNA was isopropanol-precipitated, resuspended in RNase-free H_2_O, and quantified at 260 nm by using a NanoDrop 2000 spectrophotometer (ThermoFisher Scientific Inc., Waltham, MA, USA) before microinjection. The quality and integrity of dsRNA were determined by agarose gel electrophoresis. The injection condition was set to P_i_ of 300 hpa and T_i_ of 0.3 s. Gene silencing experiments of *Duox-TLR3* RNAi and *egfp* RNAi were performed by injecting 1 μl of 2 μg/μl *dsDuox*-RNA and *dsTLR3*-RNA solution as well as *dsegfp*-RNA into the abdomen of each larva.

### Isolation of bacterial DNA from the gut and high-throughput sequencing

Total bacterial genomic DNA samples from the intestines of 27 individuals were extracted using Fast DNASPIN extraction kits (MP Biomedicals, Santa Ana, CA, USA) and stored at −20 °C prior to further analysis. The quantity and quality of extracted DNA were measured using a NanoDrop ND-1000 spectrophotometer (ThermoFisher Scientific, Waltham, MA, USA) and agarose gel electrophoresis, respectively. PCR amplification of the bacterial 16S rDNA genes at the V3–V4 regions was performed using 338F (5′ -ACTCCTACGGGAGGCAGCA-3′) and 806R (5′ -GGACTACHVGGGTWTCTAAT-3′). These primers were designed to contain a 7 nt barcode sequence for multiple samples. PCR was performed in a total reaction volume of 25 μl. The PCR conditions were as follows: initial denaturation at 95 °C for 5 min; 30 cycles of 94 °C for 20 s, 58 °C for 30 s, and 72 °C for 30 s; and a final extension at 72 °C for 2.5 min. PCR amplicons were purified with Agencourt AMPure beads (Beckman Coulter, Indianapolis, IN, USA) and quantified using the PicoGreen dsDNA assay kit (Invitrogen, Carlsbad, CA, USA). Purified PCR product libraries were quantified by Qubit dsDNA HS Assay Kit, amplicons were pooled in equal amounts, and pair-end 2 × 300 bp sequencing was performed using the Illumina Miseq PE2500 platform with MiSeq Reagent Kit version 3 (Shanghai Personal Biotechnology Co., Ltd., Shanghai, China). A total of 21 gut samples were subjected to high-throughput sequencing, including three gut samples of 8- to 10-day-old larvae with no treatment of RNAi and 18 gut samples of the *egfp* RNAi- and *Duox*-*TLR3* RNAi-treated larvae from 4, 8, and 12 days post-RNAi (DPR).

Data analysis was conducted by the bioinformatics software called Quantitative Insights into Microbial Ecology (QIIME, v. 1.8.0)[42]. We get clean sequence reads by removing of the primer sequence, truncation of sequence reads less than an average quality of 20 over a 30 bp sliding window based on the Phred algorithm, and removal of sequences that had a length of <150 bp, as well as sequences that contained mononucleotide repeats of >8 bp[26]. These strict criteria resulted in nearly 94% of the reads being retained.

FLASH was used to extend the length of short reads by overlapping paired-end reads for genome assemblies[27]. First, we sorted exactly the same sequence of clean reads according to their abundance and filtered out the singletons and used Usearch tool to cluster under 0.97 similarity. After chimera detection the remaining high-quality sequences were clustered into operational taxonomic units (OTUs) by UCLUST[28]. Selecting the highest abundance sequence from each OTU Library as the representative sequence by using default parameters. OTU taxonomic classification was conducted by BLAST search of the representative sequences set against the 16S database of known species(RDP, Http://rdp.cme.msu.edu)[29] using the best hit[30]. Reads that did not match a reference sequence at 97% identity were discarded. Bioinformatics and sequence data analyses were mainly performed using QIIME and R packages (v. 3.2.0). Using QIIME software calculates the alpha diversity index of a sample including richness and diversity indices (observed species [Sobs], Chao, abundance-based coverage estimator [ACE], and Shannon) and dissimilarity matrices (Bray–Curtis and weighted UniFrac)[31, 32].

### Statistical analysis

Results are shown as the average ± SEM of three independent biological samples. Each experiment was repeated three times. Comparison between the two independent samples were performed with student’s t-test. Multiple comparisons were conducted by one-way ANOVA and Duncan’s test using SPSS 20(IBM Corporation, Armonk, NY, USA). Significant was set at p < 0.05 The graphs were made using GraphPad Prism 5.0 (GraphPad Software, La Jolla, CA, USA).

## Results

### BSFL significantly inhibited pathogenic bacteria in the conversion of pig manure

We investigated the inhibitory action of BSF in the natural environment. Under normal pig manure conversion circumstances, BSFL significantly reduced *S. aureus* and *Salmonella* spp. counts in pig manure from day 2 to day 8. The number of *S. aureus* and *Salmonella* spp. was 5.47 and 6.71 log CFU/g on the 0 conversion day, respectively. On the 8th day after conversion, the number of *S. aureus* and *Salmonella* spp. reduced to 2.31 and 1 log CFU/g, respectively. BSFL exhibited the greatest success in reducing *S. aureus* and *Salmonella* spp. counts in pig manure on days 2, 4, and 6; suppression was also substantial on day 8 (Fig 1).

**Figure 1.**
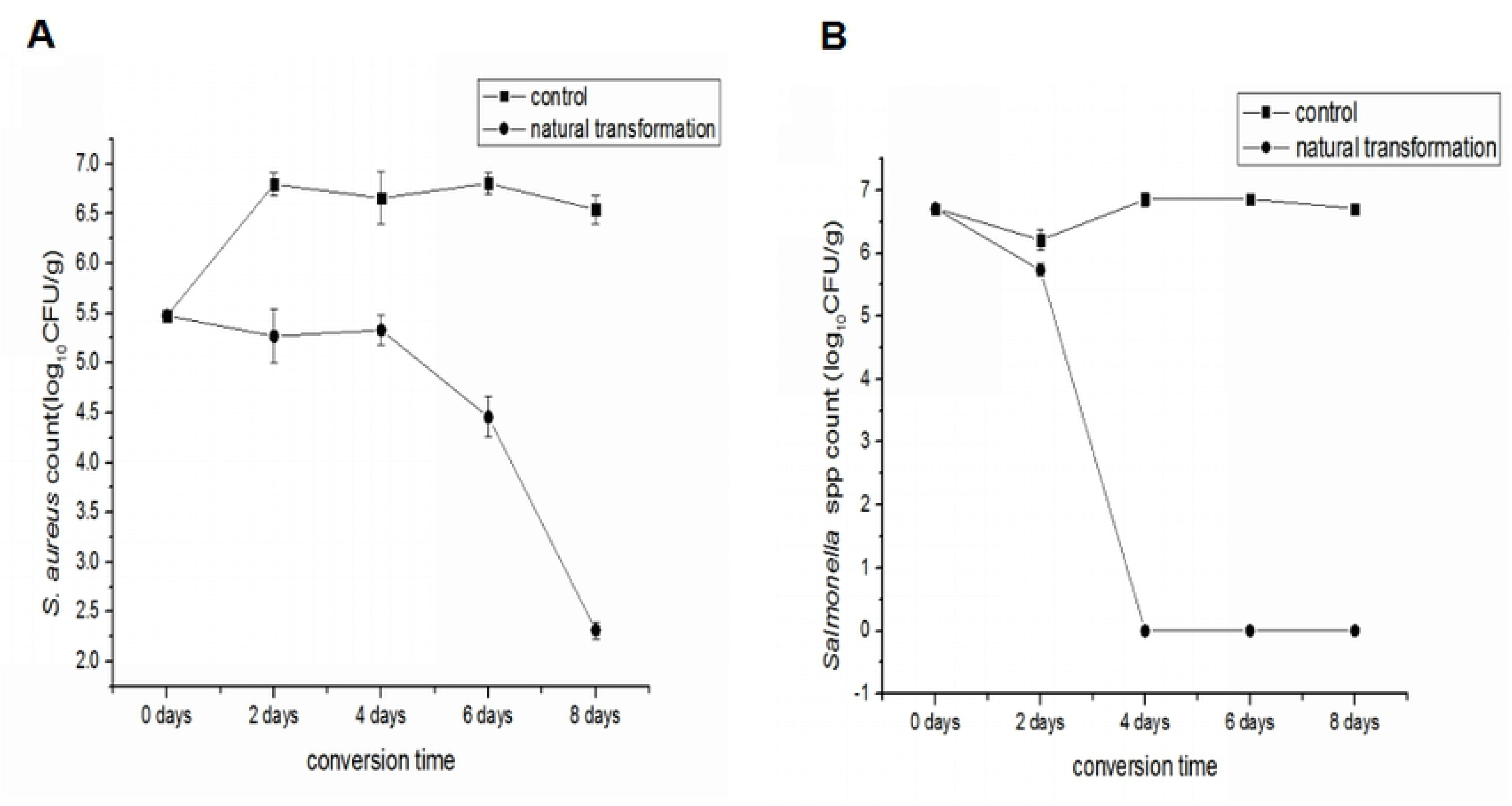
Mean *Staphylococcus aureus* (A)and *Salmonella* spp. (B) cfu/g (mean ± SEM) during 8 days transformation in pig manure with1008 days black soldier fly larvae (control) and stored at 27℃, 60%-70%RH, and a photoperiod of 16:8 (L:D) h in a growth chamber.

**Figure 2.**
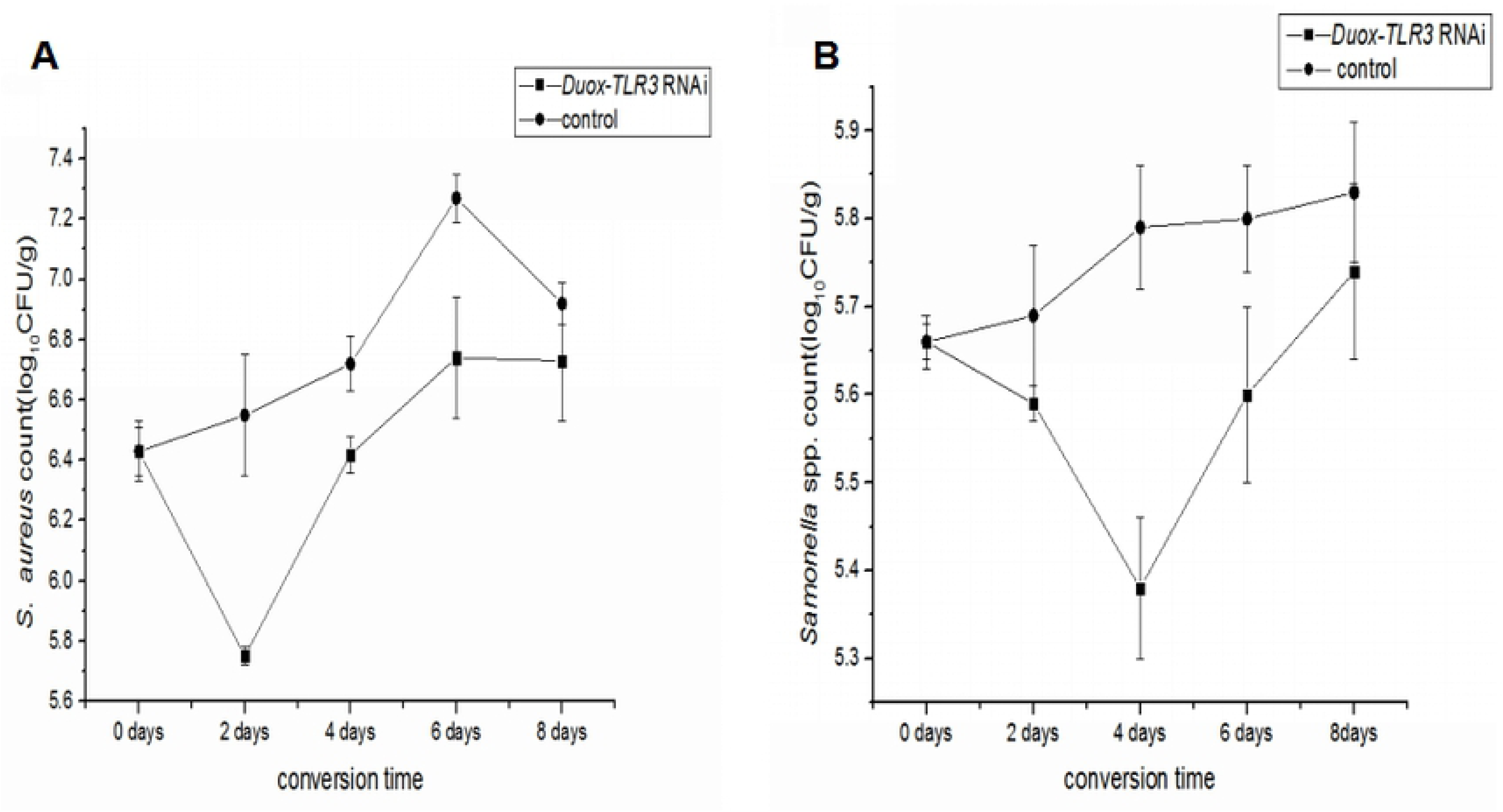
Mean *Staphylococcus aureus*(A) and *Salmonella* spp.(B). log cfu/g± SD during 8 days transformation in *Duox-TLR3* RNAi larvae group with (control) 8-d-old black soldier fly larvae and stored at 27℃, 60 –70%RH, and a photoperiod of 16:8 (L:D) h in a growth chamber.

### Attenuation of bacteriostasis after *Duox-TLR3* RNAi

After RNAi of *Duox-TLR3* in BSFL, we simulated the experiment of pig manure conversion by BSFL using the sterilized feed. First, 10^5^ CFU/g *Salmonella* spp. and 10^6^ CFU/g *S. aureus*, as well as 100 *Duox-TLR3* RNAi-injected larvae, were added to the sterilized feed. On days 0 and 8 of conversion, the *S. aureus* count was 6.43 and 6.73 log CFU/g, respectively, and the *Salmonella spp*. count was 5.66 and 5.74 log CFU/g, respectively. In the *Duox-TLR3* RNAi group, BSFL reduced the inhibition of *S. aureus* at all time points, and the interference effect was maintained for 4-8 days. Meanwhile, BSFL completely lost the inhibitory effect on *Salmonella* spp. at all points. Thus, the larvae induced a diminished effect on pathogen inhibition in Duox-TLR3 RNAi-injected larvae compared with the natural pig manure transformation conditions (Fig2).

### Sequence analysis and expression profiles of *BsfDuox* and BsfTLR3

We searched the *BsfDuox* and *BsfTLR3* cDNA from the genome database of the BSF. *BsfDuox* was 4.2 kb and encoded 1540 amino acids (S1Fig), whereas *BsfTLR3* was 1.134 kb and encoded 378 amino acids (S2 Fig). The structure of *BsfDuox* and *BsfTLR3* was similar with the previously reported *Duox* and *TLR3*of *B. dorsali*s, suggesting that they may have the same function. The *BsfDuox* gene was highly expressed in the egg; first-, second-, and fourth-instar larval; and adult stages, but it was weakly expressed in the third-instar larval stage (S5A Fig). The *BsfTLR3* gene was highly expressed in the egg; first-, second-, third-, and fourth-instar larval stages, but it was weakly expressed in the adult stage (S5B Fig). Therefore, the *BsfDuox* and *BsfTLR3* genes existed in BSF and were highly expressed in the larval stage.

### *BsfDuox* and *BsfTLR3* genes were induced upon infection

To investigate the role of *BsfDuox* and *BsfTLR3* in the immune system response, we detected *BsfDuox* and *BsfTLR3* gene expression in a gut upon oral infection with *Salmonella* spp. and *S. Aureus* induced a 4.05-times in *BsfDuox* gene expression with a peak at 12 h POI (Fig 3A). Consistent with this, ROS level was increased at 4, 12, and 24 h (Fig 3B). Thus, *BsfDuox* gene expression may be regulated by factors secreted by *S. aureus* and *Salmonella* spp. To date, uracil secreted by bacteria is the only bacterial ligand[33] that can activate *Duox* activity. Our results showed that *S. aureus* induced a 1.76-fold increase in *BsfTLR3* gene expression at 1 h POI, whereas *Salmonella* spp. induced a 1.75-fold increase in *BsfTLR3* gene expression at 4 h POI. Thus, gram-positive and gram-negative bacteria could induce *TLR3* gene expression (Fig3C).

**Figure 3.**
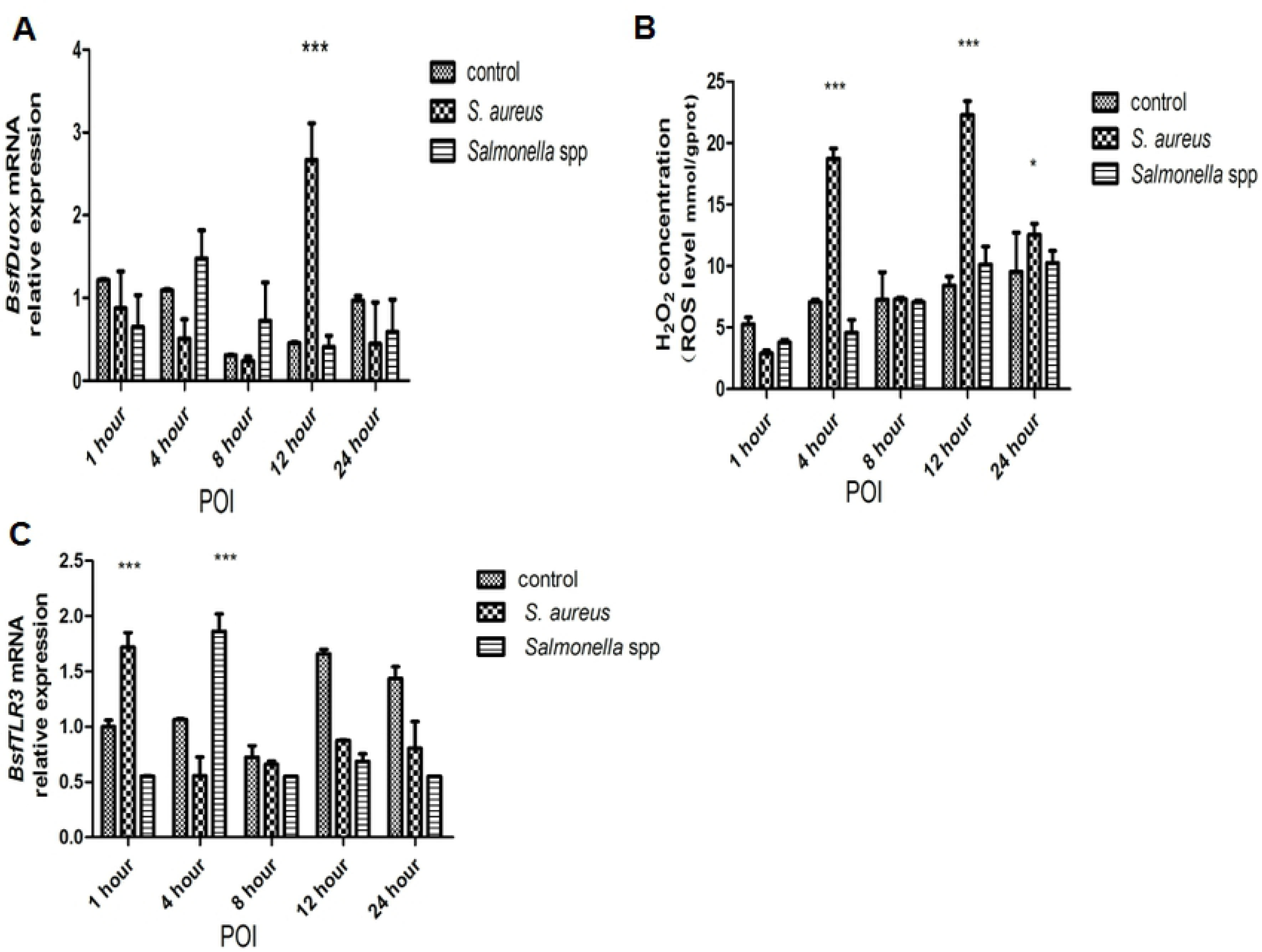
The response of the *BsfDuox* gene and *BsfTLR3* gene in the gut during oral infection. (A) Expression levels of *BsfDuox* at different time points in whole guts (without Malpighian tubules) after oral infection with *S. aureus* and *Salmonella* spp. (B)Expression levels of *BsfTLR3* at different time points in whole guts (without Malpighian tubules) after oral infection with *S. aureus* and *Salmonella* spp. (C) The total intestinal ROS levels were quantified with flies at different time points after oral infection. Data are representative of three independent experiments (mean ± SEM). Statistical comparison was based on Student’s t–test (\**p*< 0.05,\*\**p*< 0.01, \*\*\**p*< 0.001). Different letters indicate a significant difference in *BsfD*uox expression and *BsfTLR3* expression and ROS levels among the oral infection with *Salmonella* spp. or *S. aureus.*

### Sequential difference between the intestinal *Duox-ROS/Toll* signaling pathway after zoonotic pathogen challenge

We studied the temporal differentiation of the immune response of Duox-ROS immunity and Toll signaling pathway by feeding the zoonotic pathogen *S. aureus* to larvae. Infection of zoonotic pathogens could induce the immune response of the Duox-ROS system and Toll signaling pathway in a short time.

*S. aureus* induced a 2.65-fold increase in ROS in the host intestine after 4 h (Fig 3B) but a two-fold increase in *Bsfdorsal* gene in the Toll signaling pathways after 8 h (Fig 4A). The effector gene *Dif* increased by six-fold after 12 h POI compared with the control (Fig 4B). Thus, the immune response of the Duox-ROS system to zoonotic pathogen challenge was earlier than that of the Toll pathway.

**Figure 4.**
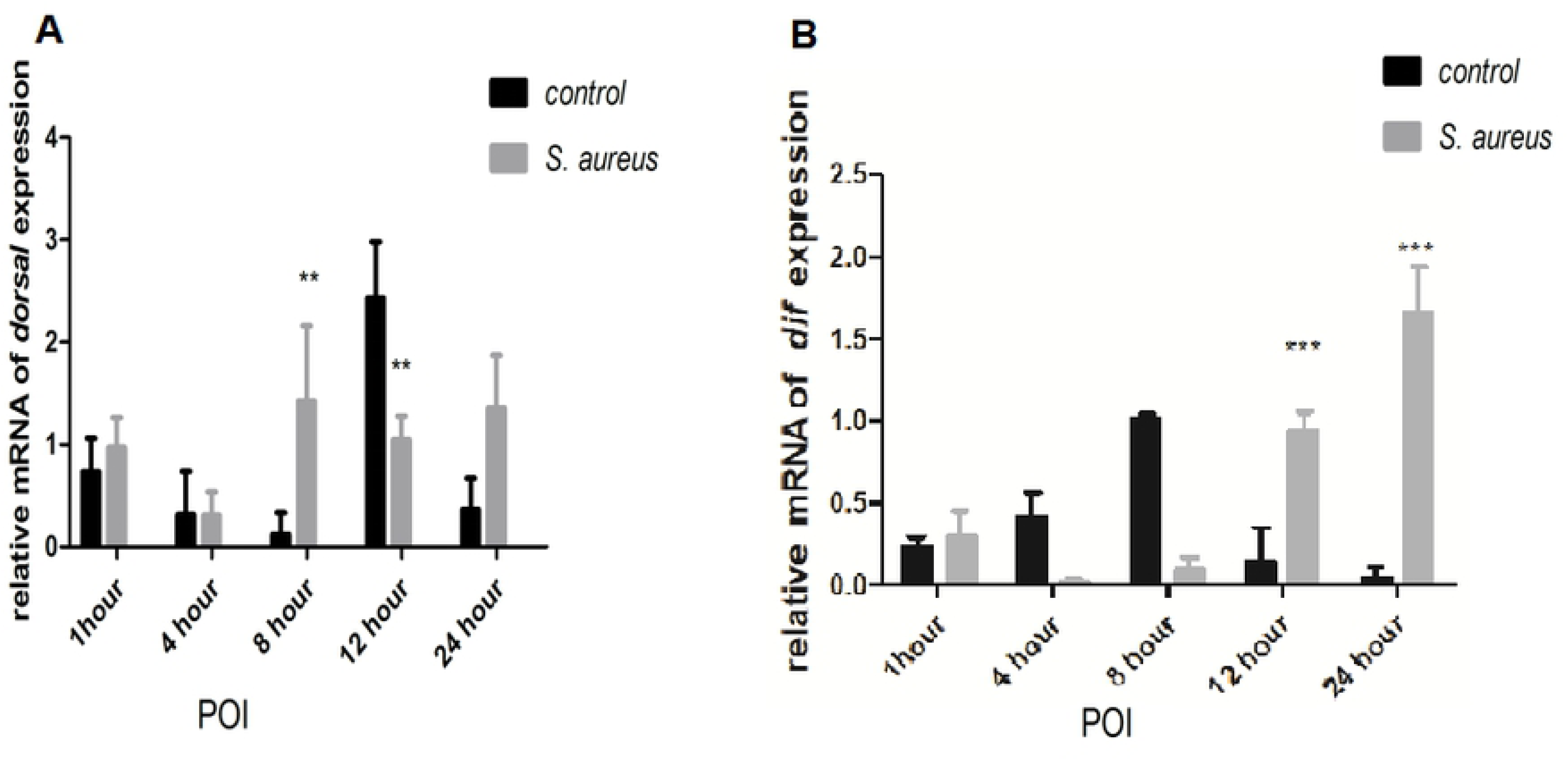
(A) The expression level of *Bsf dorsal*gene at different time points after oral infection of *S. aureus*(B) The expression level of *Bsf dif* gene at different time points after oral infection of *S. aureus.*Values are the mean ± SEM of three independent experiments. Statistical comparison was based on Student’s t–test(\**p*< 0.05,\*\**p*< 0.01, \*\*\**p*< 0.001 with Student’s t–test)

### *BsfDuox-TLR3* regulated gut bacterial density

On the basis of the above results of Figure 3, we concluded that the *BsfDuox* and *BsfTLR3* genes were activated differentially with zoonotic pathogens. Subsequently, we tested the change in composition of different intestinal symbionts, that reacted to the silencing of the *Duox-TLR3* gene by injecting larvae with *ds-Duox-TLR3*. The level of *BsfDuox* gene transcript was inhibited at 4-8 days, *BsfDuox* gene expression varied from 85% to 88% compared with the *egfp*RNAi control (Fig5A). However, *BsfDuox* expression increased in *Duox-TLR3* RNAi-treated larvae at 10 days and then returned to the basal expression level at 12 days. The level of ROS followed the same pattern with a decrease of 48%-57%as compared with the *egfp* RNAi control at 4-8 days and an increase of 21% at 10-12 days (Fig 5B). The level of *BsfTLR3* expression was inhibited at 6-10 days, *BsfTLR3* gene expression varied from 77% to 88% compared with the *egfp* RNAi control. *BsfTLR3* gene expression was 4.48 times higher in *Duox-TLR3* RNAi-treated larvae at 12 days than in the *egfp* RNAi control (Fig5C). Interestingly, the analysis of the overall bacterial density by qRT-PCR showed that *Duox-TLR3* RNAi larvae had more bacteria at 4-10 days compared with the control. The bacteria load returned to the wild-type level at 12 days (Fig 5D).

**Figure 5.**
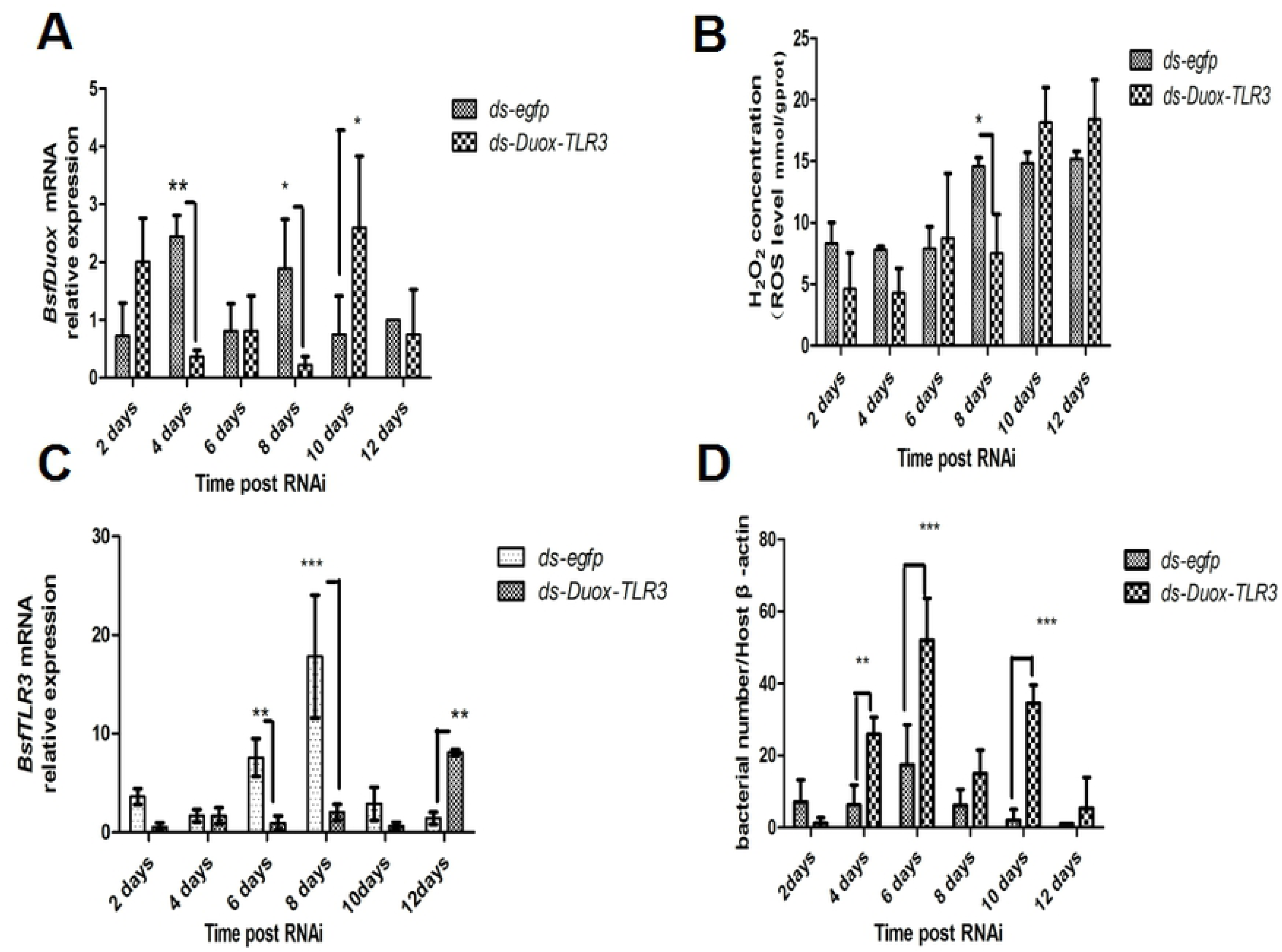
Interference effects of RNAi of the *BsfDuox*-*TLR3* gene. (A) The expression level of the *BsfDuox* gene at different time points after injecting ds-*D*uox-*TLR3* and ds-*egfp*. (B)The expression level of the Bsf*TLR3* gene at different time points after injecting ds-*D*uox-*TLR3* and ds-*egfp.* (C) The total intestinal ROS levels at different time points. (D) The total gut bacterial load at different time points. Data are representative of three independent experiments (mean ± SEM). Statistical comparison was based on Student’s t–test (\**p*<0.05, \*\**p*<0.01,\*\*\**p*< 0.001).

The AMP level of cecropin expression in *Duox-TLR3* RNAi larvae decreased by 26.14-36 times compared with that in the control at 8 and 10 days (Fig 6A). The AMP level of ubiquitin expression in *Duox-TLR3* RNAi larvae decreased by 8.71-41.55 times compared with that in the control at 6, 8, and 10 days (Fig6B). The AMP level of stomoxyn ZH1 expression in *Duox-TLR3* RNAi larvae decreased by 7.93 times compared with that in the control at 10 days (Fig6C). The results of 16s DNA sequencing of the intestinal tract of BSF showed that the beneficial symbiont bacteria in *Duox-TLR3* RNAi-injected larvae decreased by 3.4% compared with that in *egfp* RNAi-injected larvae (Fig7A), whereas the pathogenic bacteria in *Duox-TLR3* RNAi larvae increased by 44.8% compared with that in *egfp* RNAi larvae at 4-8 days post-RNAi (DPR; Fig7B). However, these differences were not statistically significant. The depletion of these two immune genes in BSFL reduced AMP and ROS production, leading to the decrease in symbiont bacteria and increase in pathogenic bacteria. Furthermore, the silencing of *BsfDuox* and *BsfTLR3* genes blocked *Duox*, *TLR3,* and ROS production, suggesting that the oral inoculation of pathobionts following RNAi could not suppress the effects of *BsfDuox* and *BsfTLR3* knockdown (S6Fig).

**Figure 6.**
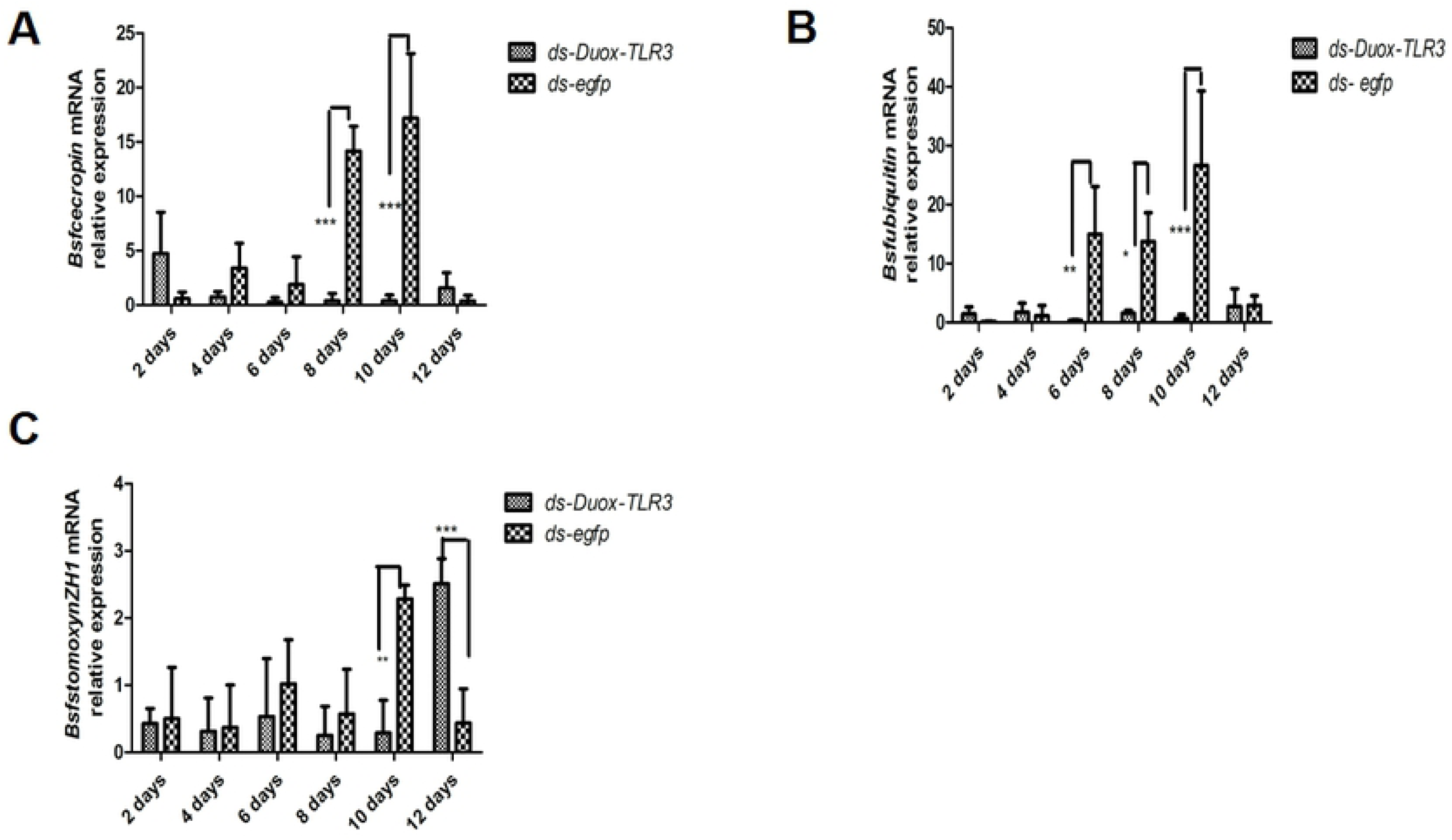
The expression level of antimicrobial peptide genes (A) *Bsf cecropin,* (B) *Bsf ubiquitin,* (C)*Bsf stomoxynZH1*, in black soldier fly larvae following the *Duox* and *TLR3* RNA interference. Data are representative of three independent experiments (mean ± SEM). Statistical comparison was based on Student’s t–test (\**p*<0.05, \*\**p*<0.01, \*\*\**p*< 0.001).

**Figure 7.**
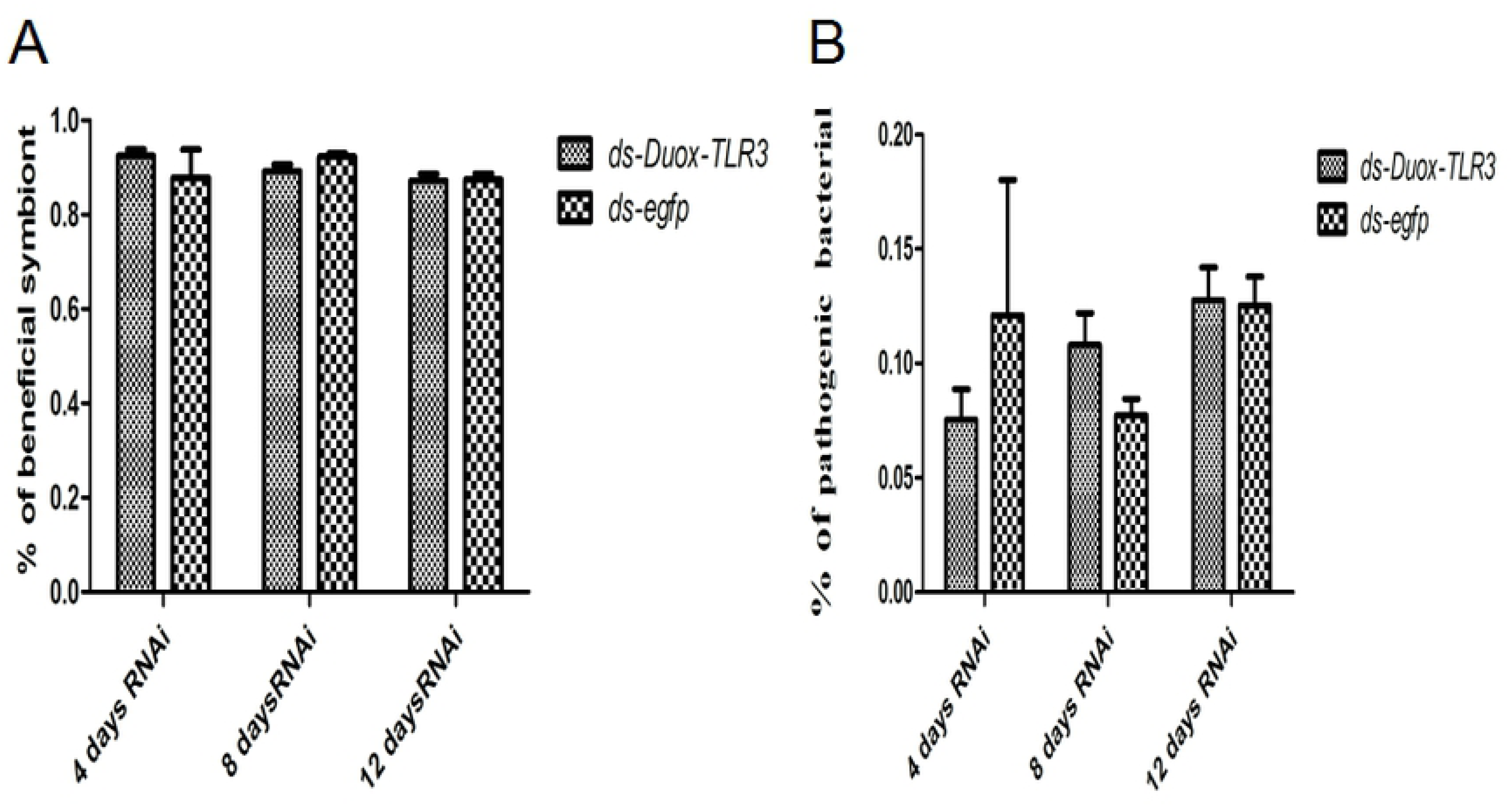
(A) The beneficial symbiont in *Duox-TLR3* RNAi larvae and *egfp* RNAi larvae at 4,8,12 days post dsRNA interfence (B)The pathogenic bacteria in *Duox-TLR3* RNAi and *egfp* RNAi larvae at 4, 8, 12 days post dsRNA interfence. Values are the mean ± SEM of three independent experiments.

The *BsfDuox-TLR3* gene by RNAi reduced the ROS and AMP levels, resulting in an increase in the number of intestinal bacteria. We attributed the low *BsfDuox* and *BsfTLR3* gene expression levels and ROS levels observed at 4-8 days to the increase in bacteria at a time when RNAi was effective. Conversely, high levels of *BsfDuox* gene expression and ROS at 10 days resulted from the increase in bacteria at a time when RNAi was ineffective. Thus, the larvae’s innate immune genes *BsfDuox* and *BsfTLR3* were actively in regulating the homeostasis of the gut bacterial and in controlling the bacterial load in the midgut, and exposure to increased pathogens resulted in the increased production of some of these anti-pathogenic factors.

### *BsfDuox-TLR3* regulated the intestinal bacterial community homeostasis

*BsfDuox* and *BsfTLR3* regulated bacterial community density but whether *BsfDuox* and *BsfTLR3* affect the gut bacterial community composition remains unknown. We then investigated the bacterial composition in *egfp* RNAi-treated and *Duox-TLR3* RNAi-treated larvae by MiSeq Illumina high-throughput sequencing. The rarefaction curves moved toward saturation representing the bacterial community well (S7A Fig). Rank abundance curve showed a rich species composition (S7B Fig). Overall, five bacterial phyla were detected in BSF samples, namely, Proteobacteria, Firmicutes, Bacteroidetes, Actinobacteria, and TM7, which composed 63.2%, 18.7 %, 16.4%, 1.29%, and 0.41% of the bacterial communities in the gut, respectively. Differences between the bacterial community of *egfp* RNAi and *Duox-TLR3* RNAi samples were calculated using the UniFrac metrics, which measures phylogenetic dissimilarities between microbial communities[34]. Genus- and species-level profiling histograms and principal coordinate analyses based on weighted UniFrac demonstrated great variation in the composition of the gut microbial community upon *Duox*-*TLR3* RNAi and *egfp* RNAi treatment, especially on days 4 and 8 post-dsRNA injection (Fig8A). Principal coordinate analysis showed separation of *Duox-TLR3*RNAi- and *egfp* RNAi-treated samples along the major component 1(pc1) and major component 2 (pc2) axes, which explained 53.07% and 13.77% of data variation (S8Fig), respectively. The analysis of the control *egfp* RNAi samples further confirmed that the intestinal bacterial composition was rich in diversity, and the major genus members of the gut community of BSF were *Providencia*,*Morganella*, *Wohlfahrtiimonas*, and *Dysgonomonas,* whereas the minor members were *Lactococcus*, *Comamonas*, *Pseudomonadaceae*, and *Leucobacter*(Fig 8A). At 4 days, *Duox-TLR3* RNAi larvae with a decrease of 12% and 14.8%abundance of *Providencia* and *Dysgonomonas*than control, respectively. However, the relative abundance of others indigenous bacteria increased by 1.17 times in the *Duox-TLR3* RNAi larvae than in the control larvae (Fig8B). The relative abundance of minor bacterial *Lactococcus* increased to detectable levels in the *Duox*-*TLR3*RNAi larvae compared with that in the control larvae (Fig8C). Thus, the larvae with reduced *BsfDuox* and *BsfTLR3* gene expression displayed a significant difference in bacterial community composition, possibly because of variations in the resistance of bacteria to ROS killing activity. At 8 days, the abundance of *Pseudomonadaceae*, a minor bacterium, in *Duox-TLR3* RNAi larvae increased significantly by twofold compared with that in the controls. The relative abundance of *Comamonas* at 8 days was similar with that at 4 days in *Duox-TLR3* RNAi larvae (Fig8E), which was due to the reduced ROS and AMP production levels. After *ds*-*Duox-TLR3* injection at 12 days, disordered intestinal bacterial communities stimulated the expression of the *BsfDuox* gene and the production of ROS to suppress non-symbionts. Finally, we observed a return to wild-type bacteria taxa composition in *Duox-TLR3* RNAi-treated larvae at 12 days, excluding *Dysgonomonas* and *Morganella*, which remained high in *Duox-TLR3* RNAi-treated larvae. Therefore, the *Duox*-*TLR3* gene had a pivotal role in regulating the structure of the bacterial community in BSFL (Fig 8F). In *Strongylocentrotus intermedius*, the petidoglycan and PolyI: C could increase the expression of *SiTLR* but LPS, ZOA could not induce the expression level of *SiTLR.* These results suggested that *SiTLR11* was functionally involved in the immune response triggered by double-stranded RNA (dsRNA) viruses and gram-positive bacteria[35]. In mosquitoes, intestinal microbiata could inhibit the development of plasmodium and other human pathogens through activation of basal immunity[9]. In the present study, the gene interference of *Duox-TLR3* resulted in a decrease in the number of beneficial symbioticbacteria *Providencia* and *Dysgonomonas* and an increase in the number of conditional pathogenic bacteria *Pseudomonadaceae*, leading to reduced resistance to pathogenic bacteria in the environment.

**Figure 8.**
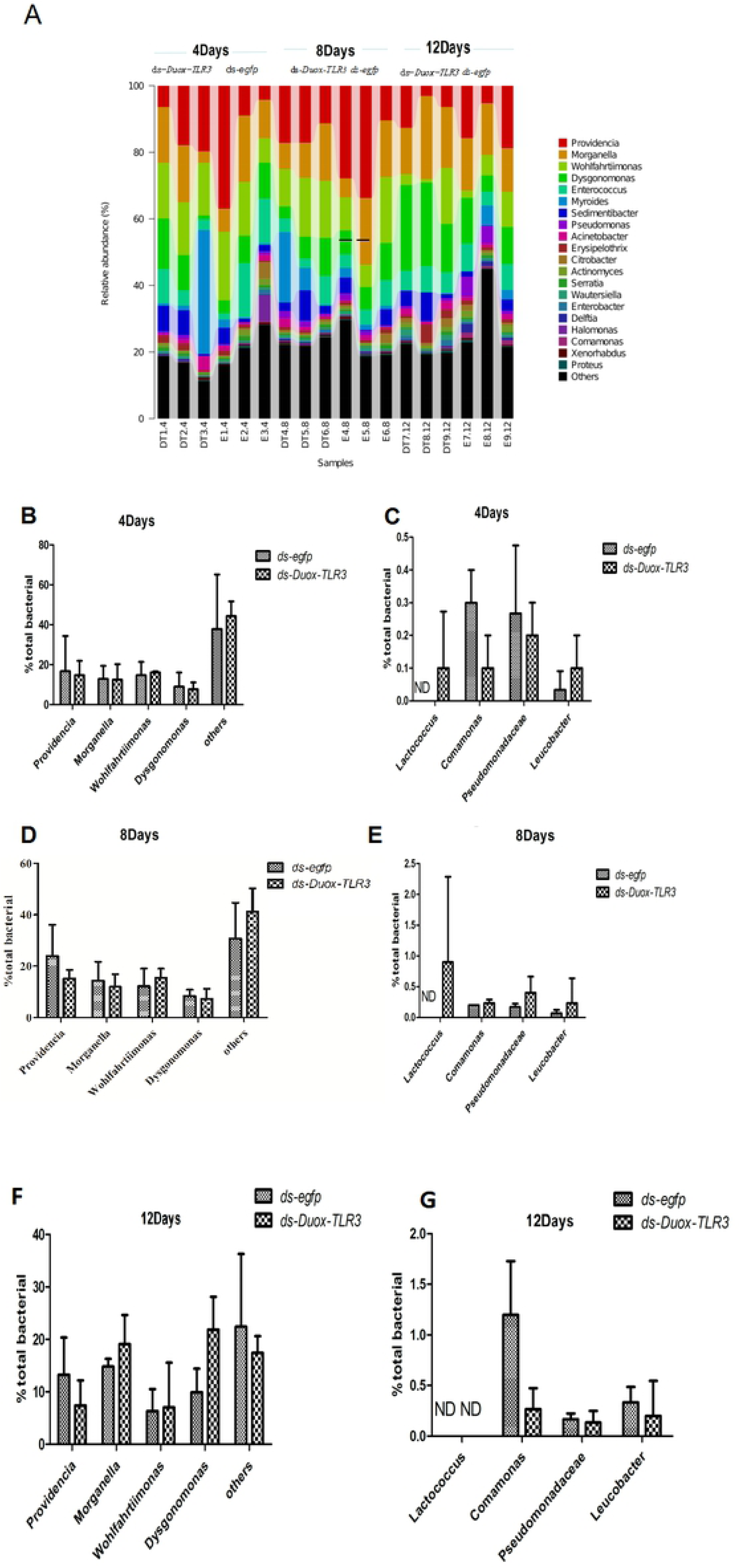
*BsfDuox-TLR3*gene regulates the composition and structure of gut bacterial community. (A) Taxonomic breakdown at the family level grouped by ds-*egfp* and ds-*Duox-TLR3* treatments. (B-G) Relative abundance of different bacterial taxa after injecting ds-*Duox-TLR3*and ds-*egfp* at 4, 8 and 12 Day. Data are representative of three independent experiments (mean+s.e.m.). Statistical comparison was based on Student’s t–test (\**p*<0.05). ND, not detected.

With the use of PICRUST, we predicted functional potentials associated with different RNAi treatments. The intestinal microbiota of *Duox-TLR3* RNAi-treated larvae was more enriched with genes involved in replication, repair, infectious diseases, carbohydrate metabolism, and amino acid metabolism but less enriched with genes involved in the biosynthesis of other secondary metabolism and enzyme family compared with that of *egfp* RNAi-treated larvae (Figs 9A and 9B).

**Figure 9.**
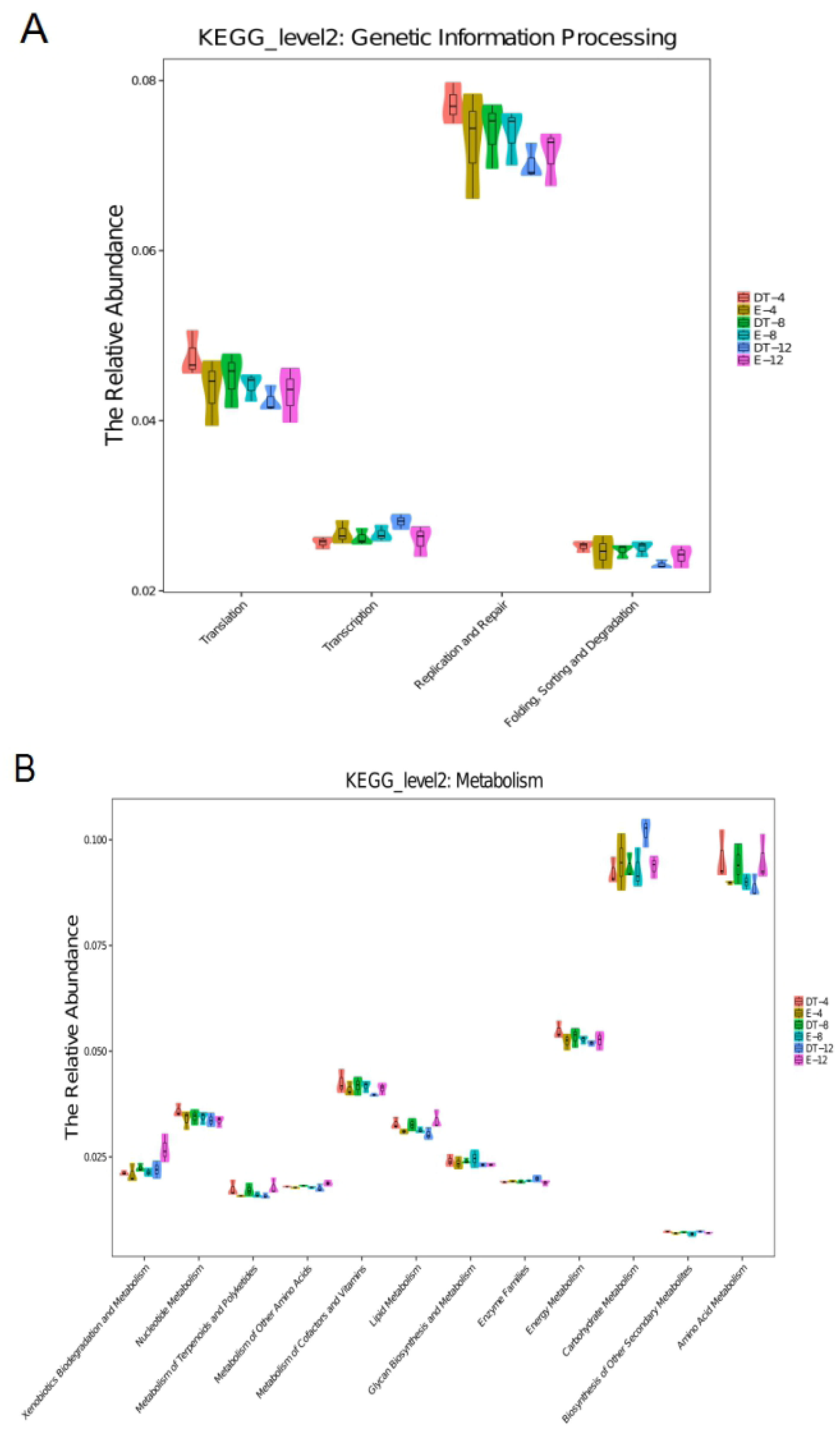
All of the predicted KEGG metabolic pathways are shown at the second hierarchical level and grouped by major functional categories. (A) KEGG_level2: Genetic information processing. (B) KEGG_level2: Matabolism.

### Microbial community richness was altered by *BsfDuox-TLR3* RNAi

We looked into the effect of *Duox*-*TLR3*RNAi on microbial diversity by a standardized approach that evaluated community richness. At 4 days, the Sobs (1192.3 ± 165.65), Chao (1598.92 ± 128.86), ACE (1628.12 ± 145.603), and Shannon (7.19 ± 0.34) indices of the intestinal microorganisms in *Duox-TLR3* RNAi larvae were the same as the Sobs (1351.3 ± 164.712), Chao (1452.85 ± 223.53), ACE (1488.46 ± 202.115), and Shannon (6.83 ± 1.07) indices in the control larvae (Fig10A). Meanwhile, the richness metrics of Chao and ACE in *Duox-TLR3* RNAi larvae significantly increased at day 8. In the control larvae, 1387 Sobs were identified, which was almost identical to those in *Duox-TLR3* RNAi larvae. The Chao, ACE, and Shannon indices increased by 31.17%, 31.6%, and 17.4%, respectively, in the *Duox-TLR3* RNAi larvae compared with those in the control larvae at day 8 (Fig10B).

Finally, no significant difference was noted in the Sobs (1316 ± 125.86 vs. 1370.66 ± 151.29), Chao (1464.42 ± 158.89 vs. 1381.29 ± 163.44), ACE (1470.208 ± 142.86 vs. 1,398.33 ± 185.09), and Shannon (7.138 ± 0.297 vs. 7.27 ± 0.517) indices between *Duox-TLR3* RNAi and *egfp* RNAi larvae at day 12 (Fig 10C). Thus, *BsfDuox-TLR3* gene silencing by RNAi led to increased bacterial diversity compared with the control, possibly because of the low solubility of ROS and AMP.

**Figure10.**
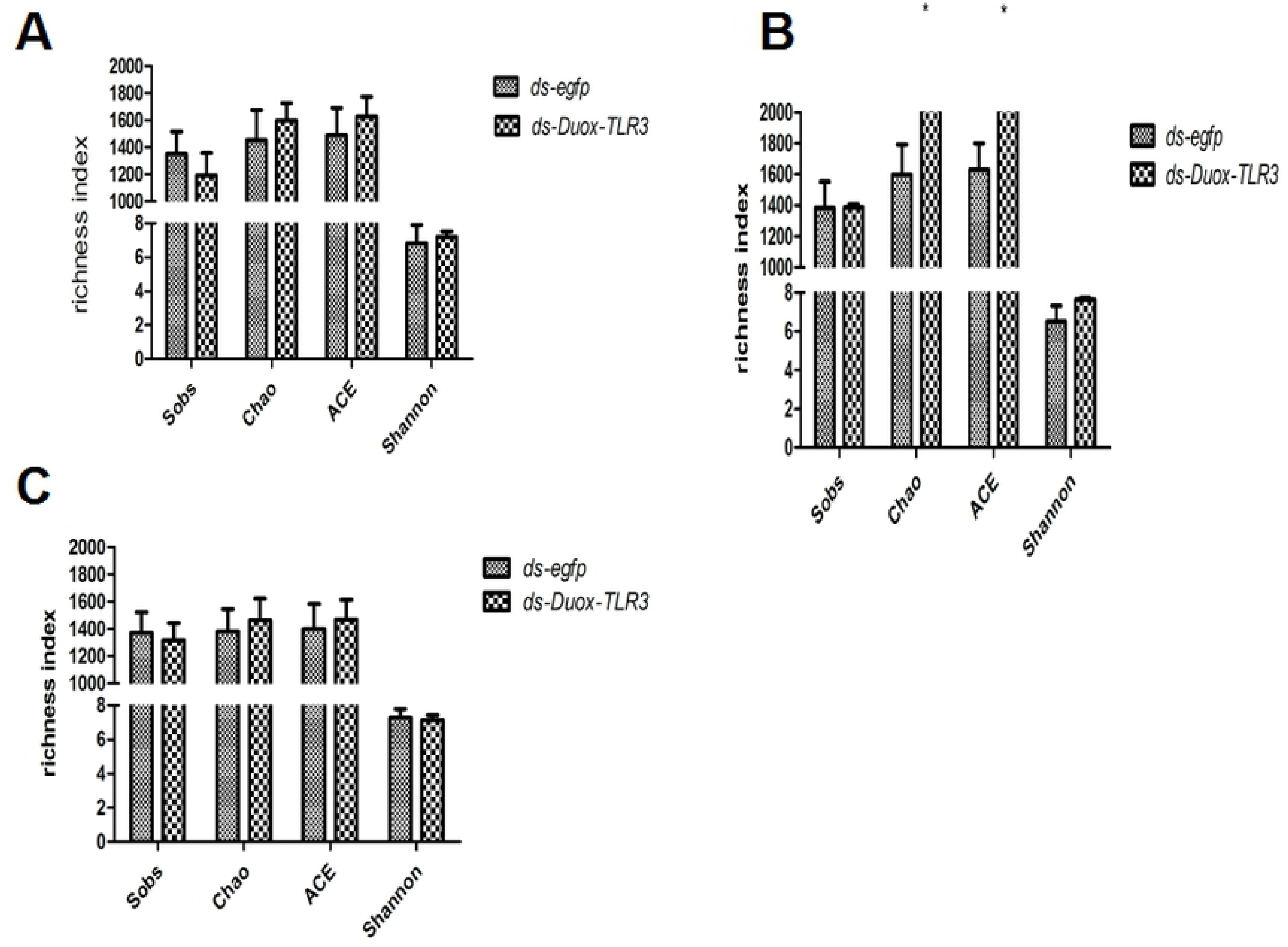
Effects of RNAi of the *BsfDuox-TLR3*gene on diversity metrics. Richness measured as observed operational taxonomic units (OTUs; Sobs), Chao, ACE and Shannon indices of gut bacterial communities from different treatment at three timepoints. (A) 4 DPR. (B) 8 DPR. (C) 12 DPR. Data are representative of three independent experiments (mean ± SEM). Statistical comparison was based on Student’s t–test (*p*).

## Discussion

Intestinal microorganisms directly or indirectly affect the immunity balance of intestinal epithelium cell, affecting the internal environment and development of the host intestine. Microbial populations in the gut regulate host immunity by balancing between an efficient immune response to inhibit foreign pathogens in the gut colonization and proliferation and an immune tolerance to the commensal microbes[36]. Conditional pathogens that activate the immune response rely mainly on AMPs and ROSthat actto restrict the growth and proliferation of invading microorganisms[37, 38]. Toll pathway-mutant larvae lacking AMP expression generally exert reduced resistance and lethal effects to gut infection. By contrast, *Duox* inactivation leads to uncontrolled of bacteria in the gut of *Drosophila*[14], illustrating that the immune system propagation of Duox-ROS system plays a major role in intestinal immune defense. *H. Illucens* has only one *Duox* gene in the genome. Structural analyses demonstrated that *BsfDuox* consists of FAD-binding and NAD-binding domains and ferric-reduct domain typical of all members of electronic delivery system responsible for H_2_O_2_ generation, as well as extracellular peroxidase homology domain (PHD). The PHD shows myeloperoxidase activity, thereby enabling the conversion of H_2_O_2_ to HOCl in the presence of chloride ion. PHD of Duox is vital for the host immune defense system(S3Fig). BSF has one *TLR3* gene, which primes the immune response. Structural analyses demonstrated that *TLR3* had three Leucine-rich repeat domain and was responsible for the protein binding and cell signaling functions (S4Fig)

With the advent of sequencing technology, many bacterial communities have been identified in the gut of various insects, including *B. dorsali*s. Indigenous bacterial community composition was of medium complexity in *B. dorsalis*, which is composed of Porphyromonadaceae, Streptococcaceae, and Sphingobacteriaceae and unclasssifed bacteria. *BdDuox* silencing led to a decreased abundance of Enterobacteriaceae and Leuconostocaceae in the gut and overgrowth of minor pathobionts[10]. Compared with the intestinal bacteria community of *B. dorsali*s, that of the BSF was relatively complex. In this study, BSF also used the Duox-ROS immune system and Toll signaling pathway as a means of intestinal immune defense. The gene expression profiles of *BsfDuox* and *BsfTLR3* at different development times indicated that they may have key function in host development. The immune defense of the *Duox* and *TLR3* genes has been studied well, but their combined role in the regulation of intestinal microbial homeostasis and bacteriostasis is rarely reported. In this study, we successfully achieved the silencing of the target gene *BsfDuox-TLR3*, which could last for 4 days. The expression level of the *BsfDuox* gene was downregulated at 4-8 days of treatment, whereas that of the *BsfTLR3* gene was downregulated at 6-10 days of treatment. The maintenance of the RNAi effect in a short time is ideal for the study of changes in intestinal microflora homeostasis. Furthermore, silencing of the *BsfDuox-TLR3* gene in larvae led to the increase in density and changes in composition and diversity of intestinal indigenous bacterial community. This finding was similar with the report on the inactivation of *Duox* and intestinal bacterial reproduction in *B.Dorsalis*[10]. The relative proportion of dominant intestinal commensal bacteria *Dysgonomonas* and *Providencia*decreased, whereas that of *Lactococcus* and *Pseudomonadaceae* increased in *Duox-TLR3* RNAi-treated larvae at 4-8 days. Thus, silencing the *BsfDuox-TLR3* gene in the intestines allowed the increase of harmful bacteria. The result may suggest that the depletion of these two immune genes in BSFL resulted in a proliferation of the minor pathogenic bacteria, which may have proven the potential decrease in anti-environmental pathogenic bacteria immune responses.

The dominant symbiotic bacteria in the gut of BSF belonged to the family of Enterobacteriaceae, particularly in the genus *Providencia*. Enterobacteriaceae is a commonly found symbiotic taxon in insects, and BSF specifically belongs to the γ-proteobacteria, a class that includes dominant symbiotic bacteria in many insect lineages[39]. Therefore, these bacterial symbionts potentially play a similar role in the biology of this fly species. Enterobacteriaceae, as dominant commensal bacteria in the intestinal tract of insects, may plays an important role in improving host fitness by preventing the colonization of foreign pathogens[40] and contribute to nitrogen fixation[41]*. Providencia* spp. are gram-negative bacteria that was one of Enterobacteriaceae family. They are resistant to two kinds of antibiotics including colistin and tigecycline, making *Providencia* multidrug-resistant. *Providencia* species which could resistant to carbapenem antibiotics are increasingly reported. Meanwhile, the genus *Dysgonomonas*is a facultative anaerobic bacteria and belongs to gram-negative coccobacilli[42]as well as is the major species in the gut microflora of BSF and *Bactrocera tau. Dysgonomonas* spp. isolated from human blood samples show antimicrobial susceptibility that directly inhibits competitors and potent pathogens from the same niche[43]. A significant decrease in *Dysgonomonas* and *Providencia* by *Duox-TLR3* gene silencing led to an adverse effect on the host. *Dysgonomonas* and *Providencia* in the gut of BSF may enhance inhibition to *Salmonella* spp. and *S. aureus,* but additional evidence is needed to verify this claim. The bacterial composition of *egfp* RNAi larvae also drastically changed at 4-12 days, probably because of the age change in the larvae. Meanwhile, *Pseudomonadaceae* is a minor component in the BSF gut and includes four pathogenic bacteria, namely, *P. Aeruginosa*[44], *Pseudomonas pertucinogena*[45], *Pseudomonas putida*[46], and *Pseudomonas fluorescens*[47]. *P. aeruginosa* is a human pathogen distributed in soil, water, air, human skin, mucous membrane, intestinal tract, and upper respiratory tract; its strains are also pathogenic to animals. The increase in *Pseudomonadaceae* bacteria caused by silencing of the *BsfDuox-TLR3* gene could cause toxicity to the host. By contrast, oriental BSF did not exhibit pathology associated with *Pseudomonadaceae*. An increased titer of this bacterium was found in the *Duox-TLR3* RNAi larvae, indicating a different function of this bacterium in the gut of resistant versus susceptible flies. Therefore, the *BsfDuox* and *BsfTLR3* genes could regulate homeostasis by ensuring the stability of symbiotic bacteria and suppressing excessive growth of minor pathogen bacteria.

The results of this study showed that oral *S. aureus* infection to BSF could induce the expression of nucleic acid transcription factors, *dorsal* and *dif*, in the Toll pathway and *Duox* in the Duox-ROS system. We also studied the signaling role of ROS in the intestinal tract by feeding different concentrations of hydrogen peroxide (0.5%– 5%) on the host. Consequently, 1%-4% H_2_O_2_ could prime the Toll signaling pathway where the intestinal *Bsfdorsal* gene mRNA level could be increased by 26.59-191.95 times (S9 Fig). However, the molecular mechanisms of specific ROS connected to the Toll signaling pathway must be further revealed.

Overall, similar to humans, BSF larval intestine harbors a natural microbiota, participating in host metabolism and provide nutrients [48]as well as in the degradation of harmful substances[49] to protect the host from adverse factors such as natural enemies, parasites. In this study, a representative of the predominant gut immune gene *Duox-TLR3* from BSF showed antimicrobial activity that directly prevented the emergence and overgrowth of minor pathobionts. The mode of protection against an encountered pathogen was possibly due to the persistent immune responses involving free radicals and antibacterial peptides. By interfering with the *Duox-TLR3* gene, the intestinal bacterial richness and composition changed (*P.putida*, and *P. fluorescens*) of BSF increased to provoke chronic inflammation under dysbiosis conditions and weakened suppression on *Salmonella* spp. and *S. aureus.* Therefore, the natural bacterial flora is crucial to maintain the stable balance of intestinal microbial communities, resisting and eliminating foreign pathogenic bacteria for insects physiological ecology such as growth and development. Understanding the role of indigenous gut residents and insect immune system contributes to the development of novel strategies against pathogens in the environment. Our results showed that *BsfDuox* and *BsfTLR3* could regulate the gut key bacteria *Providencia*and *Dysgonomonas* homeostasis to depress zoonotic pathogens. Moreover, our findings demonstrated the necessity of considering the gut symbiotic bacteria of insects during the implementation of novel pathogenic control measures.

## Acknowledgments

This work was supported by the National Natural Science Foundation of China (31770136); National Key Technology R & D Program of China (2018YFD0500203).

## Conflict of interest

The authors declare no conflict of interest.

## Author contribution

Y Q Huang design research and finalized the manuscript; Y Q Huang performed the research; Y Q Yu and S Zhuan participated part of the experiment and data analysis. Y Q Huang and M M Cai, D Hunag analyzed the data; and Y Q Huang and J K Tomberlin, L Y Zheng, Z N Yu, J B Zhang wrote the paper.

